# Efficient lentiviral transduction of different human and mouse cells

**DOI:** 10.1101/253732

**Authors:** Gang Zhang, Taihua Wang

## Abstract

**Background:** Lentiviral vectors (LVs) allowing efficient establishment of stable transgene overexpression mammalian and human cell lines are invaluable tools for genetic research. Currently, although LV transductions are broadly adopted, they are often limited due to their low titers for efficient transduction.

**Results:** Here, we described a set of optimized, efficient techniques, which could produce sufficiently high LV titers, and, provide efficient transduction of cells. According to these optimizations, most of the mammalian and human cells, both primary cells and cell lines, could be transduced successfully with high levels of transgene stable expression, including both constitutive and induced expressions.

**Conclusions:** Our data demonstrated the highly usefulness of our optimized methods. Therefore, this study provided an efficient method for most of LV transduction experiments in vitro.

## Introduction

Stable cell lines overexpressing genes of interest are powerful, indispensable tools for investigating their functions in vitro. The third generation lentivirus vectors (LVs) are broadly adopted to efficiently establish stable cell lines due to their unique characteristics, which include the ability to transduce terminally differentiated cells [1], the resistance to transcriptional silencing [2], the competence to accommodate various different promoters [3], and the self-inactivation to confirm that the transgene expressions in targeted cells are controlled solely by the internal promoters [4], and so forth.

So far, LVs are widely used for gene delivery both in vivo and in vitro. It is well known that LVs can mediate stable gene transfer into neurons in vivo [1]. In addition, LVs can be adopted to generate transgene mice and rats with both ubiquitously expressing promoters and tissue-specific promoters [3]. Furthermore, LVs can be employed to transduce somatic cells to produce induced-pluripotent stem cells [5–7]. Finally, LVs are also applied to deliver the CRISPR/Cas9 system for multiplex genome engineering and screening [8–10], etc.

However, one of the major troubles for LV transduction is due to their low titers of some LVs adopted, hence, results in low efficiencies of transduction [10]. Generally, the third-generation LV systems contain a transfer vector carrying the genes of interest, and three packaging vectors harboring Gag-Pol, Rev, and VSVG genes, respectively [11]. The working mechanism of LVs is that the Gag-Pol precursor protein is finally processed into integrase, reverse transcriptase and structural proteins during maturation. In addition, Rev interacts with a *cis*-acting element in the transfer vector enhancing export of genomic transcripts. Moreover, VSVG protein of LV envelope membrane endows LV the ability to infect a wide range of cell types, such as primary cells, stem cells, early embryos and terminally differentiated cells, etc. [1, 3, 11, 12]. When these four separate plasmids are co-transfected into human embryonic kidney (HEK) 293T cells, each cell which contains these four plasmids can produce LV particles. Previously, we co-transfected CMV-DsRed and EF1α-EGFP plasmids into HEK293 cells with Lipofectamine 2000. We found that only a small part of the transfected cells expressed both DsRed and EGFP. Whereas, a large part of cells expressed either DsRed or EGFP, respectively (unpublished data). From this phenomenon, we could infer two possible indications. The first indication is that the DNA-Lipofectamine complexes contain different DNAs. Some of them contained both CMV-DsRed and EF1α-EGFP DNAs, and the others only contained either CMV-DsRed DNA or EF1α-EGFP DNA, respectively. The second indication is, probably, during transfection, only one DNA-Lipofectamine complex can be taken into each transfected cell. Based on this hypothesis, we reasoned that, during co-transfection of several plasmids, the efficiencies of the cells containing all the plasmids are decreased with the increasing of the number of plasmids. Indeed, lacking of any plasmid of the LV systems resulted in their functional titers below detection limit [13]. This is the major barrier to obtain sufficiently high-titer LV preparations with the third generation systems.

Therefore, in this study, we made the following optimizations for LV transduction experiment design. 1) We coated 10cm dishes with 0.001% poly-L-lysine for culturing HEK293T cells during transfection, to let the cells stick to the dishes strongly, so that the cells were growing better and more resistant to detaching from the dishes [11]. 2) We employed LV systems containing the third generation, self-inactivating lentiviral transfer vectors (pWPI and pLenti/TO, respectively), and the second generation packaging vectors (for instance, pMD2.G and psPAX2) to increase the transfection efficiencies of the necessary plasmids (in total, three separate plasmids). 3) We mixed these three plasmid DNAs thoroughly before combined with Lipofectamine, to increase the number of DNA-Lipofectamine complexes, which contained these three plasmids. 4) We developed a simpler method for LV titration to instruct our further LV transduction performance. Our data demonstrated that our LV transduction experiments are sufficiently efficient. This study provided an important method for LV transduction and establishment of stable transgene cell lines in vitro.

## Materials and methods

### Construction of lentiviral transfer vectors

The construction of the lentiviral transfer vectors was described in details previously [14–17]. The vectors included pWPI/α-SynWt/Neomycin (Neo), pWPI/α-SynA30P/Neo, pWPI/α-SynA53T/Neo, pWPI/EGFP/Neo, pWPI/hPlk2Wt/Neo, pWPI/hPlk2K111M/Neo, pWPI/hPlk2T239D/Neo, pWPI/hPlk2T239V/Neo, pWPI/β5Wt/Neo, pWPI/β5T1A/Neo, pWPI/GDIWt/Neo, pWPI/GDIR218E/Neo, pWPI/GDIR240A/Neo, pWPI/Rab3AWt/Neo, pWPI/Rab3AT36N/Neo, pWPI/Rab3AQ81L/Neo, pLenti/TO/β5Wt/Puromycin (Puro), pLenti/TO/β5T1A/Puro. The bicistronic LV pWPI (Addgene plasmid 12254), which carries an EF1α-IRES-Neo cassette for the constitutive expression of dual genes driven by EF1α promoter and mediated by IRES, was modified by replacing EGFP sequence with Neo sequence, to form pWPI/Neo LV [14]. The pLenti CMV/TO Puro DEST (Addgene plasmid 17293) inducible LV, which carries a CMV/Tet-On (TO)-PGK-Puro cassette, provides the inducible expression of the genes of interests driven by CMV/TO promoter and regulated by doxycycline, and the constitutive expression of Puro driven by PGK promoter, respectively [15, 18].

### Cell culture

All animal experiments were performed according to the guidelines established in the Canadian Guide for the Care and Use of Laboratory Animals. HEK293, HEK293T, HEK293/TO, SHSY5Y (ATCC), SHSY5Y/TO, adult mouse whole brain neural progenitor cells (NPCs), and BV-2 mouse microglial cells were thawed from liquid nitrogen. HEK293, HEK293/TO, and HEK293T cells were cultured in Dulbecco’s modified eagle medium (DMEM) containing 10% fetal bovine serum (FBS) in 10-cm dishes. SHSY5Y and SHSY5Y/TO cells were cultured in DMEM plus 15% FBS in 10-cm dishes (Wisent Biocenter). NPCs were cultured in the medium DMEM/F12/N2 (Gibco) containingβFGF (20ng/ml, Invitrogen), EGF (20ng/ml, Invitrogen) and heparin (5μg/ml, Sigma), in T75 flasks [19, 20]. BV-2 mouse microglial cells were cultured in RPMI1640 (Gibco) plus 10% FBS, respectively [21].

### Production of lentiviruses

HEK293T cells were thawed from liquid nitrogen and cultured in DMEM containing 10% FBS, growing for a week in 10cm dishes, passage at least three times. 10cm dishes were coated with 0.001% poly-L-lysine (Sigma) for 15 minutes at room temperature (RT), then the poly-L-lysine was removed thoroughly by siphoning [11]. When HEK293T cells were passaged, at first, the cells were washed with Ca^2+^, Mg^2+^-free DPBS twice (Wisent Biocenter), then, treated with 0.25% Trypsin-EDTA (Wisent Biocenter) for 1 min at RT, and then pipetted up and down about five times in order to separate almost all of them into single cells. To obtain even distribution of the cells across the surface of the plates, the cells were mixed thoroughly by pipetting up and down with 10ml serological pipettes before seeding, after seeding the cells, do not rock the dishes, or more cells would move to the central parts of the dishes, and decrease the following transfection efficiencies. The HEK293T cells were cultured until about 70-80% confluence, and the lentiviral transfer vectors and packaging vectors, psPAX2 (Addgene plasmid #12260) and pMD2.G (Addgene plasmid #12259), were co-transfected into HEK293T cells, respectively, by Lipofectamine 2000 according to the manufacturer’s instructions (Invitrogen). Importantly, to gain efficient production of LVs, the DNA amounts of transfer vectors and packaging vectors were adjusted according to their sizes, and let their molecular ratios were about 1:1:1, respectively. For each transfection, the DNAs of the transfer vector and two packaging vectors were added into 1ml DMEM (without FBS) according to the supplementary data table 1, and the DNA mixtures were mixed thoroughly by low-speed vortex. And then, 40μl Lipofectamine 2000 was diluted in 1ml DMEM (without FBS), mixed gently by inversion of the tube for about 10 times, incubated for 5min at RT. After 5min incubation, the diluted DNAs and diluted Lipofectamine 2000 were combined, mixed gently by inversion the tube for about 10 times, and incubated for 20 min at RT to form Lipofectamine-DNA complexes [14]. Finally, the 2ml complexes were added to each well containing HEK293T cells and 8ml DMEM + 10% FBS medium, mixed gently by rocking the plates. The cells were incubated at 37°C in a CO_2_ incubator overnight. According to different purposes, different media were used to harvest the lentiviruses. For HEK293, SHSY5Y and BV-2 cell infection, 6ml fresh DMEM + 2% FBS medium was used [11]. For NPCs infection, NPC culture medium (DMEM/F12/N2 + 20ng/mlβFGF + 20ng/ml EGF + 5μg/ml heparin) was used for harvesting the viruses. 48 hours post-transfection, the supernatants were harvested for LVs. The viruses were centrifuged at 2500 x g for 10 min, and filtered through 0.45μm filters. The viruses were used for titration and infection freshly, or stored at −80°C freezer.

**Table 1.** The relations between the LV titers and the sizes of the genes of interest.

| Vector name | Inserts | Sizes (kb) | Titers (Mean ± SD) |
| --- | --- | --- | --- |
| pWPI/Neo | NO | 0 | $4.18 \times 10^7 \pm 9.58 \times 10^6$ |
| pWPI/α-SynWt | α-SynWt | 0.64 | $2.55 \times 10^7 \pm 5.83 \times 10^6$ |
| pWPI/α-SynA30P | α-SynA30P | 0.64 | $4.55 \times 10^7 \pm 3.12 \times 10^7$ |
| pWPI/α-SynA53T | α-SynA53T | 0.64 | $4.58 \times 10^7 \pm 9.04 \times 10^6$ |
| pWPI/EGFP | EGFP | 0.75 | $3.33 \times 10^6 \pm 3.85 \times 10^6$ |
| pWPI/Rab3AWt | Rab3AWt | 0.78 | $3.93 \times 10^7 \pm 3.86 \times 10^6$ |
| pWPI/β5Wt | β5Wt | 0.9 | $3.11 \times 10^7 \pm 1.49 \times 10^7$ |
| pWPI/β5T1A | β5T1A | 0.9 | $1.6 \times 10^7 \pm 5.14 \times 10^6$ |
| pWPI/GDIR218E | GDIR218E | 1.44 | $3.26 \times 10^7 \pm 8.22 \times 10^6$ |
| pWPI/GDIR240A | GDIR240A | 1.44 | $2.44 \times 10^7 \pm 2.25 \times 10^6$ |
| pWPI/hPlk2Wt | hPlk2Wt | 2.1 | $4.96 \times 10^6 \pm 133 \times 10^6$ |
| pWPI/hPlk2K111M | hPlk2K111M | 2.1 | $6.92 \times 10^6 \pm 287 \times 10^6$ |
| pWPI/hPlk2T239D | hPlk2T239D | 2.1 | $3.55 \times 10^6 \pm 198 \times 10^6$ |
| pWPI/hPlk2T239V | hPlk2T239V | 2.1 | $3.04 \times 10^6 \pm 2.07 \times 10^6$ |
| pLenti/TO/β5Wt/Puro | β5Wt | 0.9 | $1.63 \times 10^6 \pm 5.66 \times 10^5$ |
| pLenti/TO/β5T1A/Puro | β5T1A | 0.9 | $5.04 \times 10^4 \pm 2.33 \times 10^4$ |

### Titration of lentiviruses

The titration was performed with 6-well plates. First of all, 6-well plates were coated with 0.001% poly-L-lysine (Sigma) for 15 min at RT [11]. Then HEK293 cells were split, and seeded in each well. When the cells were about 70-80% confluent, the medium was removed. Then 900μl fresh DMEM + 2% FBS medium were added into each well, respectively. The viruses were thawed on ice, and 6-well plates were used to make 10 x fold dilutions (Figure 1). After the viruses were thawed, in the first well, 100μl virus was added into 900μl DMEM +2% FBS medium to make 10 times dilution, mix them sufficiently by shaking the plate. Then, in the second well, 100μl 10 x fold diluted virus was added into 900μl medium to make 100 times dilution, and repeated on until to make 10^7^ or 10^8^ dilutions. The infected cells were incubated at 37°C in a CO_2_ incubator overnight. The medium with viruses was replenished with 2ml fresh DMEM + 10% FBS medium. 48 hours after infection, add 400-800μg/ml G418 (BioShop) gradually to select infected HEK293 cells with Neomycin expression [22], and 1.5μg/ml Puro for infected HEK293 cells with pLenti/TO/β5Wt/Puro and pLenti/TO/β5T1A/Puro LVs [23, 24]. After two weeks of selection, the live cell colonies were directly counted under microscope or stained with crystal violet (Sigma). Briefly, 1% crystal violet solution was prepared in 10% ethanol, then, the medium of the selected cells was removed, and the cells were washed twice with DPBS (Wisent Biocenter). Then, 1ml crystal violet solution was added into each well, and incubated for 10 min at RT. And then, the staining solution was removed and the cells were washed twice with DPBS. Finally, the stained cell colonies were counted with naked eyes. The titers of the LVs were calculated as below: Titers (TU/ml) = colony number x dilution factors (such as 10^4^, 10^5^, etc.) x (1000/900) (if not the last well) to adjust into 1 ml volume (Figure 2, 3).

**Figure 1.**
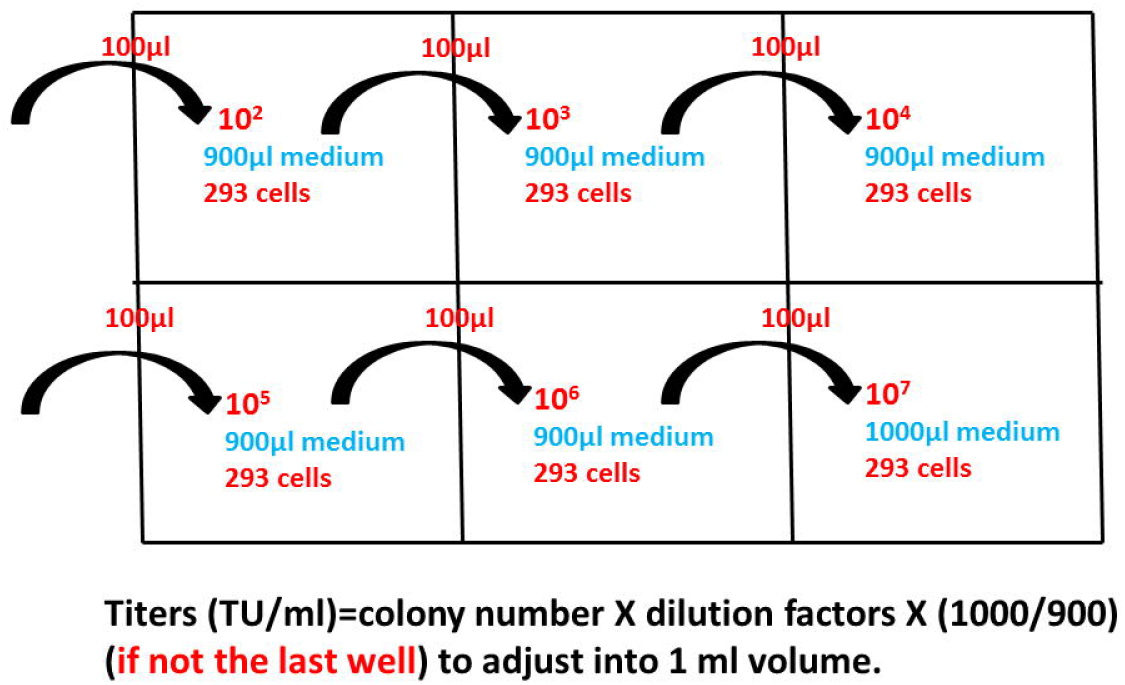
The scheme of titration using 6-well plates. 6-well plates were used for titration. The plates were coated with 0.001% poly-L-lysine for 15 min at RT. Then HEK293 cells were seeded in each well and let cells grow up to about 70-80% confluent. And then, each well was replenished with 900μl fresh DMEM + 2% FBS medium, respectively. The viruses were thawed, and made 10 x fold dilutions directly in the 6-well plates with cells. In each well, 100μl diluted virus was added into 900μl medium, and repeated on until 10^7^ or 10^8^ dilutions. **PLEASE noted** that, in order to make 10^7^ or 10^8^ dilutions, the early 10 fold dilutions were made using another 6-well plate, and did not show in this scheme. The infected cells were selected with 400-800μg/ml G418 for neomycin expression, and 1.5μg/ml puromycin for pLenti/TO/Puro vector infection. After two weeks of selection, the live cell colonies were directly counted under microscope or stained with crystal violet. Finally, the stained cell colonies were counted with naked eyes. The titers of the lentiviruses were calculated as indicated in the figure.

**Figure 2.**
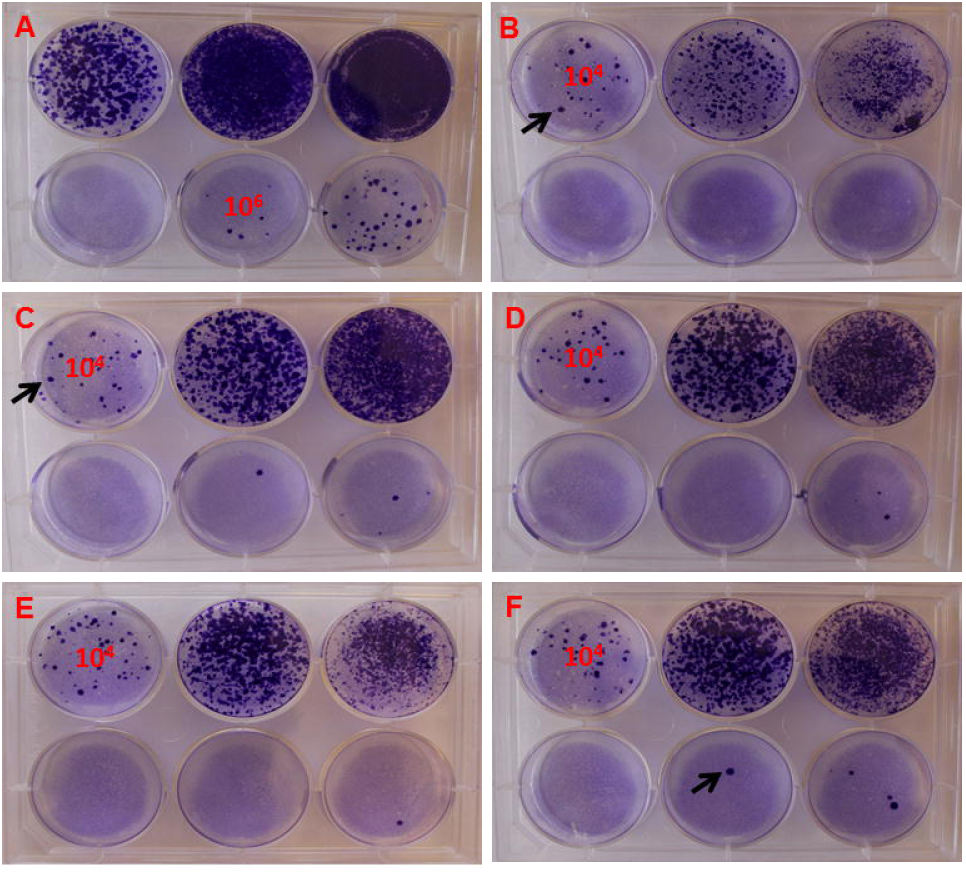
The titration of pWPI/Neo empty vectors and vectors carrying EGFP, hPlk2Wt and K111M, T239D, T239V mutants, stained with crystal violet (n=1). The big blue-stained colonies (indicated with black arrows) represent several infected cells grew together to form a big cell colony, but each of them was counted as one colony for calculating the titers. A. pWPI/Neo, 10^6^ dilution well was chosen for counting the live cell colonies; B: pWPI/EGFP/Neo; C: pWPI/hPlk2Wt/Neo; D: pWPI/hPlk2K111M/Neo; E: pWPI/hPlk2T239D/Neo; F: pWPI/hPlk2T239V/Neo. For B, C, D, E and F, the 10^4^ dilution wells were chosen for counting the live cell colonies for calculating their titers.

**Figure 3.**
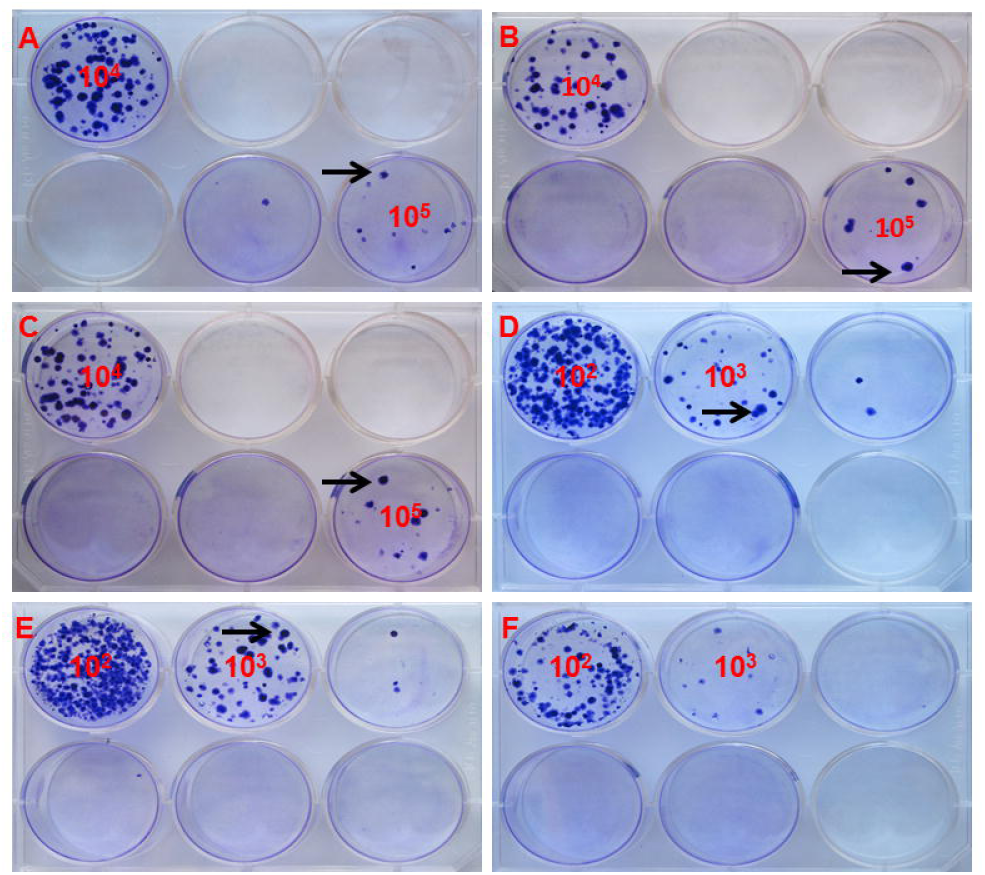
The titration of pLenti/TO/ 5Wt/Puro and pLenti/TO/ 5T1A/Puro stained with crystal violet (n=3). The big blue-stained colonies (indicated with black arrows) represent several infected cells grew together to form a big cell colony, but each of them was counted as one colony for calculating the titers. A, B, and C: pLenti/TO/β5Wt/Puro, 10^5^ wells were chosen for counting the live cell colonies; D, E, and F: pLenti/TO/β5T1A/Puro, the 10^3^ wells were chosen for counting the live cell colonies.

## Infection and selection of stable cell lines

### HEK293, SHSY5Y cells

Upon infection, the LVs were thawed on ice from −80°C freezer, and HEK293, SHSY5Y, SHSY5Y/TO cells were cultured and passaged as described above. The 10-cm dishes were coated with 0.001% Poly-L-lysine for infection. When the cells were about 80% confluent, the medium was removed thoroughly, and 6ml LV supernatants were directly added into the 10-cm dishes, respectively. The cells were infected for 6 hours or overnight. Then, the virus medium was replenished with DMEM + 10%FBS and DMEM +15%FBS, respectively. For HEK293 cells infected with pWPI/Neo vectors with respective insert genes, 500-800μg/ml G418 was added gradually to select stable cell lines for two weeks [22]. For SHSY5Y cells infected with pWPI/Neo vectors with insert genes, 300-600μg/ml G418 was added gradually to select stable cell lines for two weeks [25].

### NPCs

NPCs were cultured in T75 flasks to form neurospheres. NPC neurospheres were collected into 15ml Falcon tubes, and centrifuged at 800 RPM for 5 min. Then, the medium was removed, and washed with Ca2+, Mg2+-free DPBS twice, each time, remove the DPBS with centrifugation. And then, add 1ml Accutase (Life Technologies) to dissociate the neurospheres for 2-3 min at RT. Finally, 2ml NPC culture medium (DMEM/F12/N2 + 20ng/ml bFGF + 20ng/ml EGF + 5μg/ml heparin) was added to stop the Accutase digestion, then, pipetting up and down about five times to separate the neurospheres into single NPCs. Finally, remove the supernatant by centrifugation. 6ml pWPI/Neo lentiviruses with different genes of interest were directly added into the single NPCs, mixed gently by pipetting, and the NPCs were moved into T75 flasks, incubated overnight for infection. 48 hours after infection, 50-200μg/ml G418 was gradually added to select the stable cell lines for two weeks [26, 27].

### BV-2 cells

Upon infection, in two 15ml Falcon tubes, 1 X 10^6^ BV-2 cells were harvested into each of them, and centrifuged at 1000 RPM for 5 min, remove the supernatant. In one tube, add 5ml pWPI/EGFP/Neo virus (the titer was about 2 x 10^5^ TU/ml) to infect the BV-2 cells with the MOI (multiplicity of infection) about 1:1. In another tube, 0.4ml pWPI/EGFP/Neo virus (the titer was about 4 x 10^6^ TU/ml) to infect the BV-2 cells with the MOI about 1.6:1. The cells were mixed by pipetting gently, and incubated at 37°C in a CO_2_ incubator overnight. Then, the cells were replenished with fresh RPMI1640 + 10% FBS, and moved into 6cm dishes. 300-500μg/ml G418 was gradually used to select stable cell lines for two weeks [28].

### SHSY5Y/TO cells

For SHSY5Y/TO cells infected with pLenti/TO/β5Wt/Puro and pLenti/TO/β5T1A/Puro inducible LVs, 6-cm dishes were coated with 0.001% Poly-L-lysine, and 1.5 x 10^6^ cells were seeded each in dish. 6 hours after seeding, the medium was removed, and 6ml LVs of each were added into the dishes. The titers of pLenti/TO/β5Wt/Puro and pLenti/TO/β5T1A/Puro were 1.63 x 10^6^ ± 5.66 x 10^5^ and 5.04 x 10^4^ ± 2.33 x 10^4^, respectively. Therefore, the MOI of pLenti/TO/β5Wt/Puro virus to SHSY5Y/TO cells was 6.5:1, and the MOI of pLenti/TO/β5T1A/Puro virus to the cells was 0.2:1.

### Induction of β5Wt and β5T1A expression in SHSY5Y/TO cells

1μg/ml Doxycycline was added into the culture medium to induce the expression of βWt and βT1A mutant expression in HEK293/TO and SHSY5Y/TO cells [18].

### Western blotting

Cell lysates were prepared in cell lysis buffer (50mM Tris, pH8.3, 150mM NaCl, 1% NP-40, 1% Sodium deoxycholate, and 1 X Protease inhibitor cocktail (Sigma)). Briefly, cell culture medium was removed from 10-cm dishes, and washed with ice-cold DPBS twice. 1ml lysis buffer was added into each dish, incubated on ice for 10 min. The lysates were aliquoted into 300μl/tube. BCA Protein Assay (Thermo Scientific) was performed to test the protein concentrations according to the manufacturer’s guide. 75μl 5 X Laemmli buffer (312.5mM Tris-HCl, pH6.8, 10% SDS, 250mM DTT, 50% Glycerol, and 0.05% Bromophenol blue) was added into each tube, mixed and heated at 95°C for 10 min. Finally, the samples were stored at −80°C until running SDS-PAGE.

12%, 1.0mm SDS-PAGE gels were run to separate the sample proteins. 10μl of BenchMark^TM^ prestained protein ladder (Invitrogen), and 25-40μg of samples were loaded. Non-infected cell lysate was used as negative controls for the transgene expression. The gels were run in 1XTris-glycine SDS buffer (Bioshop). The proteins were transferred onto nitrocellulose and processed for Western blotting. Protein bands were visualized with ECL-plus Western blotting detection system (GE Healthcare), and quantified with a Storm 860 fluorescent imager (Molecular Dynamics).

Antibodies used in this study included: mouse anti-tetraHIS primary antibody (1:1000 dilution) (Qiagen), mouse anti-glyceraldehyde 3-phosphate dehydrogenase (GAPDH) primary antibody (1:1000 dilution) (Sigma). The goat anti-mouse secondary antibody was Immunopure^®^ goat anti-mouse IgG, H+L, Peroxidase conjugated, purchased from Thermo Scientific. Goat anti-mouse secondary antibody was 1:2500 diluted for probing tetraHIS, and GAPDH expression [14]. All the antibodies were diluted in Tris buffered saline (TBS, Wisent Biocenter) with 0.05% Tween-20 and 5% milk.

### Fluorescence microscopy

A Axiovert 100TV (Zeiss) fluorescence microscope equipped with a AttoArc^TM^ HBO 100W (Carl Zeiss) lamp power supply was used to observe the stable expression of EGFP in BV-2 cells after G418 selection for two weeks. Microscopic images were photographed using a digital camera (Kodak DC290 ZOOM).

### Data Statistics

Data were analyzed by Mean ± SD (standard deviation) and Student’s T-TEST.

## Results

### Titration of LVs

The titration of the LVs was performed in 6-well plates manifested in Figure 1. After two weeks of selection with G418 or puro, the live cell colonies were either directly counted under microscope, or stained with 1% crystal violet, and counted with naked eyes. The titers of the LVs were analyzed with Mean ± SD and T-TEST. The detailed TTEST data were listed in supplementary data Table 2. Generally, our data showed that the titers of LVs in this study were decreased with the increase of the sizes of the genes of interest (Figure 4, A, Table 1).

**Figure 4.**
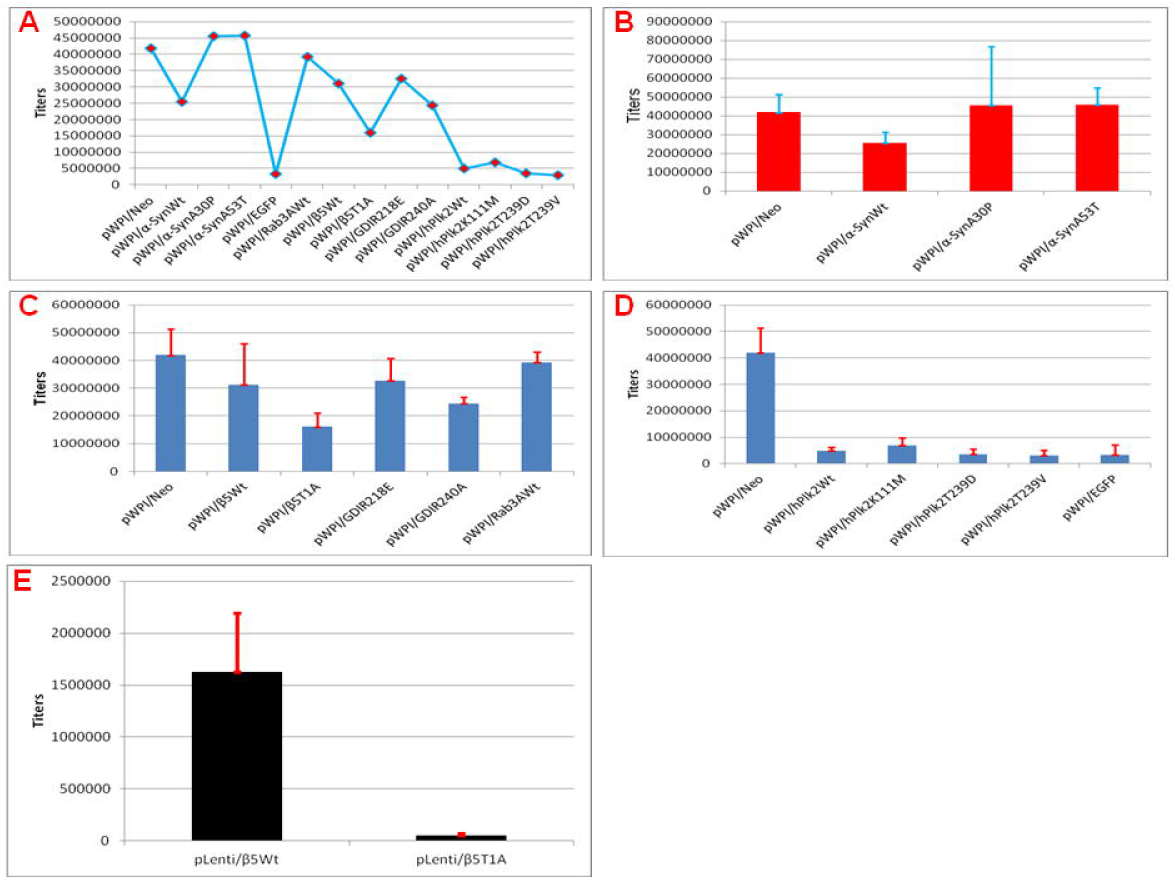
Titers of the LVs (n=3) The different lentiviruses were indicated in the figures A, B, C, D and E. The titers were analyzed by Mean ± SD and student’s T-TEST. The analyses were performed with Microsoft Windows 7, Excel software.

### Titers of pWPI/α-SynWt/Neo and mutants LVs

The titers of pWPI/Neo and pWPI/α-SynWt/Neo, pWPI/α-SynA30P/Neo, and pWPI/α-SynA53T/Neo LVs were 4.18 x 10^7^ ± 9.58 x 10^6^, 2.55 x 10^7^ ± 5.83 x 10^6^, 4.55 x 10^7^ ± 3.12 x 10^7^, and 4.58 x 10^7^ ± 9.04 x 10^6^, respectively. Compared with the titer of pWPI/Neo LV, the titers of pWPI/α-SynWt/Neo, pWPI/α-SynA30P/Neo, pWPI/α-SynA53T/Neo LVs had no significant difference (P>0.05). But the titers between pWPI/α-SynWt/Neo and pWPI/α-SynA53T/Neo LVs had significant difference (P<0.05, Figure 4. B).

### Titers of pWPI/β5Wt/Neo and pWPI/β5T1A/Neo LVs

The titers of pWPI/β5Wt/Neo and pWPI/β5T1A/Neo were 3.11 x 10^7^ ± 1.49 x 10^7^ and 1.6 x 10^7^ ± 5.14 x 10^6^, respectively. There was no significant difference between the titers of pWPI/Neo and pWPI/β5Wt/Neo LVs (P>0.05). But, the titers of pWPI/Neo and pWPI/β5T1A/Neo LVs had significant difference (P<0.05). Furthermore, the titers between pWPI/β5Wt/Neo and pWPI/β5T1A/Neo LVs had no significant difference (P>0.05, Figure 4. C).

### Titers of pWPI/GDIR218E/Neo and pWPI/GDIR240A/Neo LVs

The titers of pWPI/GDIR218E/Neo and pWPI/GDIR240A/Neo LVs were 3.26 x 10^7^ ± 8.22 x 10^6^ and 2.44 x 10^7^ ± 2.25 x 10^6^, respectively. Compared with the titer of pWPI/Neo LV, the titers of pWPI/GDI218E/Neo and pWPI/GDI240A/Neo LVs had no significant difference (P>0.05). In addition, the titers between pWPI/GDIR218E/Neo and pWPI/GDIR240A/Neo LVs also had no significant difference (P>0.05, Figure 4. C).

### Titers of pWPI/Rab3AWt/Neo and pWPI/EGFP/Neo LVs

The titers of pWPI/Rab3AWt/Neo and pWPI/EGFP/Neo vectors were 3.93 x 10^7^ ± 3.86 x 10^6^ and 3.33 x 10^6^ ± 3.85 x 10^6^, respectively. The titers between pWPI/Neo and pWPI/Rab3AWt/Neo LVs had no significant difference (P>0.05). But, the titers between pWPI/Neo and pWPI/EGFP/Neo LVs had significant difference (P<0.05). Notably, the titers between pWPI/EGFP/Neo and pWPI/Rab3AWt/Neo LVs had highly significant difference (P<0.01, Figure 4. C, D).

### Titers of pWPI/hPlk2Wt and mutants LVs

The titers of pWPI/hPlk2Wt/Neo, pWPI/hPlk2K111M/Neo, pWPI/hPlk2T239D/Neo, and pWPI/hPlk2T239V/Neo LVs were 4.96 x 10^6^ ± 1.33 x10^6^, 6.92 x 10^6^ ± 2.87 x 10^6^, 3.55 x 10^6^ ± 1.98 x 10^6^, and 3.04 x 10^6^ ± 2.07 x 10^6^, respectively. Compared with the titer of pWPI/Neo LV, the titers of pWPI/hPlk2Wt/Neo, pWPI/hPlk2K111M/Neo, pWPI/hPlk2T239D/Neo, and pWPI/hPlk2T239V/Neo LVs had significant difference (P<0.05). But the titers between pWPI/hPlk2Wt/Neo, pWPI/hPlk2K111M/Neo, pWPI/hPlk2T239D/Neo, and pWPI/hPlk2T239V/Neo LVs had no significant difference (P>0.05, Figure 4. D).

### Titers of pLenti/TO/β5Wt/Puro and pLenti/TO/β5T1A/Puro LVs

The titers of pLenti/TO/β5Wt/Puro and pLenti/TO/β5T1A/Puro LVs were 1.63 x 10^6^ ± 5.66 x 10^5^ and 5.04 x 10^4^ ± 2.33 x 10^4^, respectively. After T-TEST analysis, the titers between them were significantly different (P<0.05). When the titers between pWPI/β5Wt/Neo, pWPI/β5T1A/Neo, pLenti/TO/β5Wt/Puro, and pLenti/TO/β5T1A/Puro LVs were compared, respectively, the titers of pLenti/TO/β5Wt/Puro and pWPI/β5Wt/Neo LVs had no significant difference (P>0.05). In addition, the titers between pLenti/TO/β5T1A/Puro and pWPI/β5Wt/Neo LVs also had no significant difference (P>0.05). Furthermore, the titers between pLenti/TO/β5Wt/Puro and pWPI/βT1A/Neo LVs had significant difference (P<0.05). Moreover, the titers between pLenti/TO/β5T1A/Puro and pWPI/β5T1A/Neo LVs also had significant difference (P<0.05, Figure 4. C, E).

## Establishment of stable cell lines

HEK293, SHSY5Y, SHSY5Y/TO, NPCs and BV-2 glial cells were infected with LVs, and selected with G418 or Puro antibiotics for two weeks by gradually increasing the drug concentrations, respectively. Then, the stable transgene cell lines were tested by Western Blotting or Fluorescent Microscopy, respectively.

### Stable transgene expression in HEK293 and BV-2 cell lines

To test the transgene expression of the stable cell lines, Western Blotting and Fluorescent Microscopy were performed for their expressions in HEK293, and BV-2 cells, respectively. The open reading frames (ORF) of β5Wt and β5T1A mutant contained 873 base pairs (bp) with fused c-Myc and 6xHis tag sequences at their C-termini before the stop codon. The molecular weights of their full length precursor protein were about 31.7 kD. The mature β5 subunit was generated through two proteolytic events [29]. One was just before the active site Thr1, which was lost in β5T1A mutant, and the other was in the middle of the propeptide, which was still present in β5T1A mutant. Therefore, a mature β5T1A mutant subunit was with an extended 2.1kDa N-terminal sequence resided upstream from the T1A mutation site [30]. As a result, the mature β5Wt was about 24.7kD, whereas, the mature β5T1A was about 26.8kD (Figure 6. A, B, C, D, E and F; Supplementary data Figure 3. A and C). The ORFs of Rab3A Wt, and T36N, Q81L mutants contained 681bp with fused 6xHis tags at their N-termini immediately after start codon. Their protein molecular weights were about 25.8kD (Supplementary data Figure 3. A). The ORFs of GDI Wt and R218E, R240A mutants contained 1362bp with fused 6xHis tags at their N-termini immediately behind the start codon. Their protein molecular weights were about 51.3kD (Supplementary data Figure 3. B). Their expressions were tested with Western blotting for the 6xHis tag expression. EGFP stable expression in BV-2 cell line was observed by fluorescence microscopy (Figure 5. A, B, C and D). Our data demonstrated that all the genes described above were efficiently expressed in HEK293 and BV-2 cell lines. Some of the expression data of these genes in SHSY5Y, NPCs were published elsewhere [19, 31].

**Figure 5.**
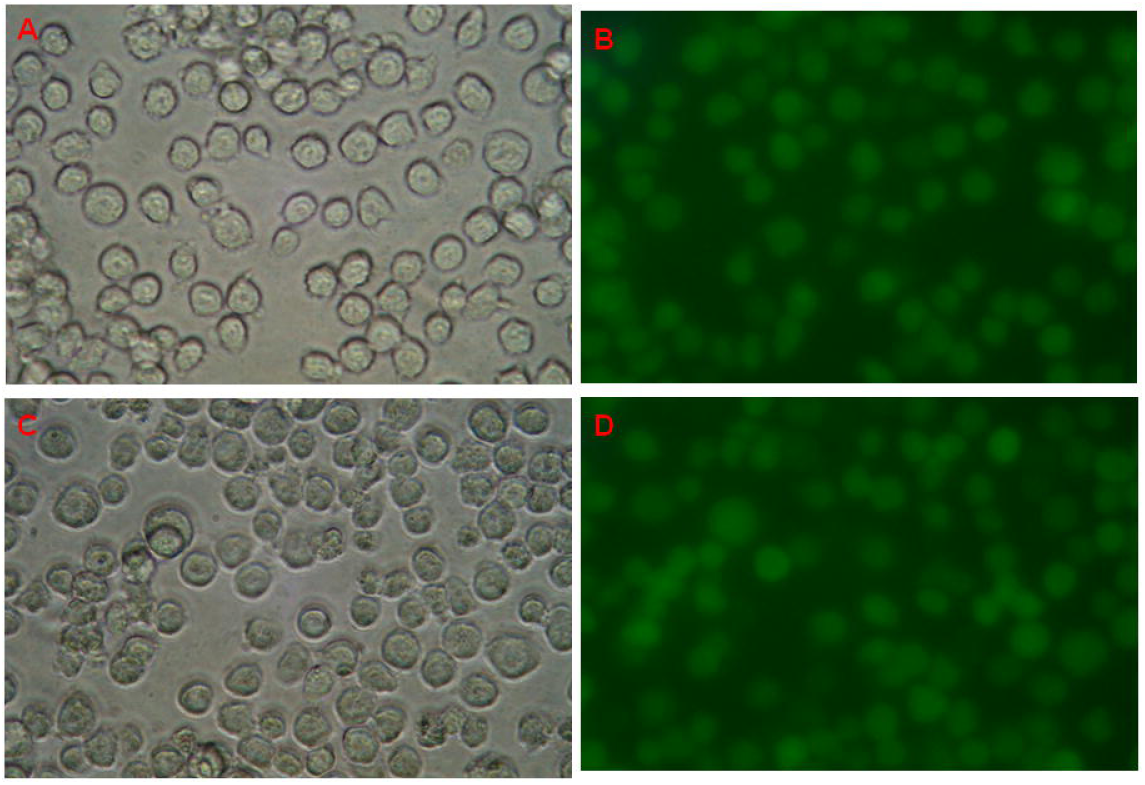
Stable expression of EGFP in BV-2 mouse microglial cells. BV-2 cells were infected with pWPI/EGFP/Neo LV and selected by G418 for two weeks to establish stable BV-2/EGFP cell lines. Under a fluorescence microscope, the same microscopic field (at 20X original magnification) was documented by digital camera images taken with grayscale (A and C) and green fluorescence (B and D). The MOI (viruses : cells) of A and B was 1.6:1, and the MOI of C and D was 1:1.

### Induced expression of β5Wt and β5T1A in SHSY5Y/TO and HEK293/TO cells

The induced expressions of β5Wt and β5T1A in SHSY5Y/TO cell lines were regulated by adding different amount of Doxycycline (Dox) in cell culture medium to induce the expression for 1 to 8 days. Their expressions were tested by Western Blotting with Tetra-His and GAPDH primary antibodies (Figure 6. A, B, C, D, E, and F), our data showed that the induced long-term stable expressions of β5Wt and β5T1A mutant in SHSY5Y/TO cells were successfully achieved. Interestingly, compared with the induced transient expression of β5Wt and β5T1A mutant in HEK293/TO cells, more mature β5Wt and β5T1A proteins were detected after induction in SHSY5Y/TO stable cell lines via LV transduction (Figure 6. A, C and E). Whereas, more full length precursor β5Wt and β5T1A proteins were detected 48 hours after induction through lipofectamine 2000 transfection in HEK293/TO cells (Supplementary data Figure 3. C).

**Figure 6.**
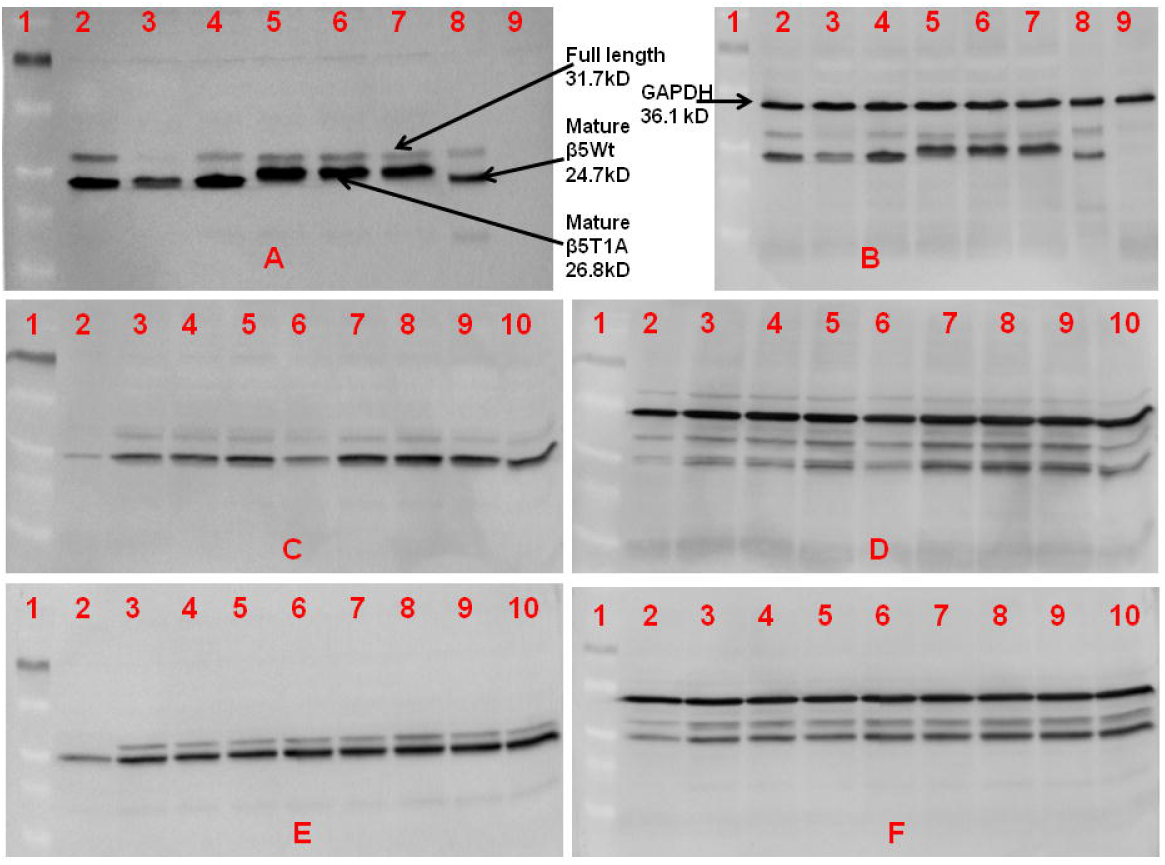
Induced expressions of β5Wt and β5T1A in SHSY5Y/TO cell lines. A and B: SHSY5Y/TO cells were infected with pLenti/TO/β5Wt and pLenti/TO/β5T1A LVs, selected with puromycin, and induced for expression by Dox for 4 days. About 40μg samples were loaded in 1.0mm, 12% SDSPAGE gel. A was probed by TetraHis primary antibody, whereas, B was tested by GAPDH primary antibody. Lane 1: Protein ladder; lane 2: β5Wt (1μg/ml Dox); lane 3: β5Wt (No Dox, not induced); lane 4: β5Wt (1μg/ml Dox); lane 5: β5T1A (1μg/ml Dox); lane 6: β5T1A (No Dox); lane 7: β5T1A (1μg/ml Dox); lane 8: 293/pWPI/β5Wt/Neo (Positive control); lane 9: SHSY5Y/TO cells (Not transduced, negative control). Lanes 2, 3, 4, and lanes 5, 6, 7 were cell lines infected respectively, and lane 3 and lane 6 were leak expression without induction. C and D: Induced expression of β5Wt in SHSY5Y/TO cells transduced by pLenti/TO/β5Wt/Puro LV, selected with puromycin, with 1μg/ml Dox for different days. About 30μg samples were loaded in 1.0mm, 12% SDS-PAGE gel. C was probed by TetraHis primary antibody, and D was tested by GAPDH primary antibody. Lane 1: protein ladder; lane 2: Not induced; lane 3: day 1; lane 4: day 2; lane 5: day 3; lane 6: day 4; lane 7: day 5; lane 8: day 6; lane 9: day 7; lane 10: day 8. E and F: Induced expression of β5T1A in SHSY5Y/TO cells transduced by pLenti/TO/β5T1A/Puro LV, selected with puromycin, with 1μg/ml Dox for different days. About 30μg samples were loaded in 1.0mm, 12% SDS-PAGE gel. E was probed by TetraHis primary antibody, and F was tested by GAPDH primary antibody. Lane 1: protein ladder; lane 2: Not induced; lane 3: day 1; lane 4: day 2; lane 5: day 3; lane 6: day 4; lane 7: day 5; lane 8: day 6; lane 9: day 7; lane 10: day 8.

## Discussion

LVs with sufficient titers are the prerequisite for efficient LV transduction. Due to the mechanisms of the third generation LVs, the titers of LVs are mainly depended on the following four factors. 1), the transfection efficiencies affect the LV titers radically. Generally, for the same LV backbones and the same produce cell lines with the same methods, the bigger the sizes of the inserts, the lower the efficiencies of the transfections. Therefore, the titers of LVs harboring longer inserts are lower than those with shorter inserts (Figure 4. A). 2), the LV systems employed are also an important factor for their titers. For example, the packaging vectors of the third generation LVs contain three separate plasmids carrying the necessary genes for assembling LV particles. Whereas, the packaging vectors of the second generation LVs contain two separate plasmids. Therefore, if the third generation lentiviral transfer vectors, such as pWPI/Neo, and pLenti/TO/Puro, together with the three separate third-generation lentiviral packaging vectors, are employed to co-transfect the HEK293T produce cells, only the individual cells containing these four vectors, simultaneously, can produce LV particles [11]. As a contrast, if the third generation lentiviral transfer vectors, together with the two separate second generation lentiviral packaging vectors, are adopted, each cell carrying these three vectors can produce LV particles. Indeed, omission of any one of these plasmids (transfer vector, envelope vector and packaging vectors, respectively) during production resulted in no detectable functional titers (which refer to the concentration of vector particles capable of transducing target cells) [13, 32]. For this reason, using the third generation transfer vector plus the second generation packaging vectors can increase the production of LVs particles, and hence, improve the efficiencies of LV transduction for the targeting cells. 3), the toxicity of the genes carried by the LV transfer vectors also can affect the LV production significantly. For instance, in our experiments, we found that the cells transfected with LV vectors carrying β5T1A mutant gene resulted in more cell death than β5Wt gene (unpublished data). As a result, the titers of pWPI/β5Wt/Neo and pLenti/TO/β5Wt/Puro are higher than pWPI/β5T1A/Neo and pLenti/TO/β5T1A/Puro in our experiments, particularly between pLenti/TO/β5Wt/Puro and pLenti/TO/β5T1A/Puro (Table 1, Figure 3, Figure 4. C). 4), the toxicity of the genes can also decrease the LV titers during the titration step. As aforementioned, when HEK293 cells were infected with LVs carrying β5Wt and β5T1A, such as pLenti/TO/β5Wt/Puro and pLenti/TO/β5T1A/Puro LVs, more infected cells with pLenti/TO/β5T1A/Puro LV were died. These died cells certainly cannot form cell colonies after two-week antibiotic selection. Therefore, the calculated titers were decreased significantly than pLenti/TO/β5Wt/Puro LVs (Figure 3, Figure 4. E, Table 1, P<0.05). Although their titers are significantly different between pLenti/TO/β5Wt/Puro and pLenti/TO/β5T1A/Puro LVs, their induced expressions could be detested efficiently in SHSY5Y/TO cells (Figure 6. A, C and E). This result further indicated that, in this instance, the calculated titers of pLenti/TO/β5T1A/Puro LV did not reveal its true number of infectious particles.

Because the self-inactivating of the third generation LVs prevented subsequent viral replication in the targeted cells [4, 11, 33], based on the above analyses, besides the inherent factors, such as the toxicity, and specific sequences of the genes of interest (for example, β5T1A and EGFP, etc, EGFP is a well-known neutral reporter in mammalian cells [34], but, the titers are significantly lower than other genes with similar sizes, Table 1), the transfection procedure is the sole controllable step to affect the LV titers. To improve the production of LVs, we optimized the cell culture and transfection protocols. Because HEK293T cells can not stick to the dish bottom strongly, some cells are easy to become floating, and decrease the transfection efficiencies. To prevent the cells to detach the dish, we coated the 10-cm dishes with 0.001% Poly-L-lysine during the LV production step [11]. In addition, when we perform the co-transfection step, we let the HEK293T cells at about 70-80% confluence, and the cells still have room for another division (293T cell line lost the contact inhibition ability). At this point, the cells are exponentially growing, so maximal transfection efficiencies might be achieved [35]. Furthermore, as aforementioned, when several plasmids were co-transfected, only a small proportion of cells contained all the plasmids. Previously, we combined CMV-DsRed plasmid with Lipofectamine 2000 to form DsRed-DNA-Lipofectamine complexes. At the same time, we also combined EF1α-EGFP plasmid with Lipofectamine 2000 to form EGFP-DNA-Lipofectamine complexes. Then, we mixed the two complexes and added into cell culture medium to transfect the cells. Surprisingly, no cells expressing both DsRed and EGFP simultaneously were found (unpublished data). This phenomenon further indicated that during transfection procedure by Lipofectamine, probably, only one DNA-Lipofectamine complex can enter into each targeting cell. Moreover, to improve the efficiencies of DNA-Lipofectamine complex formation containing the lentiviral transfer vectors and two packaging vectors, simultaneously, as many as possible, we mixed these three vectors (their molecular ratios are about 1:1:1) thoroughly by low-speed vertex. We found these optimizations could increase the productivity of LV particles. Finally, we successfully established dozens of stable transgene cell lines which efficiently expressed the genes of interest in target cells (Figure 5, 6, Supplementary data Figure 3, A and B).

There are mainly three different methods for LVs titration. One of them is called RNA titer, which is the assessment of LV RNA in supernatant [32, 36]. Usually, RNA titers can not accurately reflect the functional titer of LVs, and are substantially higher than the functional titers, which refer to the concentrations of LV particles capable of transducing target cells [32, 37–39]. Another method is designated as DNA titer, which is the measurement of LV DNA in transduced cells [32]. The third method is called functional titer (also as biological titer, BT = TU/μl, transducing units/μl) [11], in which the LV DNA expression in targeted cells is measured by staining or fluorescence expression followed by fluorescent activated cell sorting [32, 40]. Some investigators argued that this method could not distinguish cells with single or multiple integrations [13, 32]. Our data suggested that almost all the transduced cells were with nearly identically integrated copy number after LV infection in BV-2 cell lines (Figure 5, B and D). This phenomenon indicated that, during titration, in high dilution wells of infection, such as 10^3^ (Figure 3, D, E and F), 10^4^ (Figure 2, B, C, D, E and F), 10^5^ (Figure 3, A, B and C) and 10^6^ (Figure 2, A) in different experiments, the integration numbers of targeted cells might be identical, and it might be one copy of each targeted cell (Supplementary data Figure 1 and Figure 2). In our experiments, we did see some “huge” colonies (Figure 2 and Figure 3, black arrows), which were formed from several closely neighbored separate infected cells. We counted each of these “huge” colonies as one colony, and this is one of the reasons that, in our titration method, the functional titers are always lower than the real LV particle numbers. By vividly seeing the different transduced cells in different dilution wells, our titration method is very instructive for further infection of targeting cells. Finally, compared with other formats, such as using 24-well plates [11], using 100μl virus volume to make the 10 X fold dilution can radically decrease the deviations of dilution as well (Figure 1).

In this study, both constitutive and induced expressions of genes of interest are achieved efficiently in different targeting cells (Figure 5 and 6; Supplementary data Figure 3). Interestingly, the expressions of β5Wt and β5T1A in stable cell lines and in transient cells are different. In stable cell lines, both constitutive and induced expressions, more mature β5Wt and β5T1A expressions were detected than their full length precursors (Figure 6, A, C and E; Supplementary data Figure 3, A). Whereas, in transient expression cells, more full length precursors of β5Wt and β5T1A were detected than their mature forms (Supplementary data Figure 3, C). Our data demonstrated that the in vitro overexpression models are successfully established. This study provided an efficient, simple method for LV transduction of different human and mouse cells.

## Abbreviations

LVs: Lentiviral vectors
HEK293T: Human embryonic kidney 293T;
Neo: Neomycin;
Puro: Puromycin;
TO: Tet-On;
NPCs: Neural progenitor cells;
DMEM: Dulbecco’s modified eagle medium;
FBS: Fetal bovine serum;
RT: Room temperature;
MOI: Multiplicity of infection;
GAPDH: Glyceraldehyde 3-phosphate dehydrogenase;
TBS: Tris buffered saline;
SD: Standard deviation;
ORF: Open reading frames;
bp: base pairs;
Dox: Doxycycline.

## Acknowledgements

We acknowledge Dr. Didier Trono for the original pWPI vector. We thank Robert Strome for the modified pWPI/Neo/BamH I vector. We acknowledge Dr. Stanley B. Prusiner for the original pTetO-HGMoPrP vector. We thank Dr. Eric Campeau for the original pLenti CMV/TO Puro DEST vector.

## Availability of data and materials

The full sequence information of the original vectors is available in the Addgene repository (www.addgene.org): pWPI (Addgene plasmid 12254), pLenti CMV/TO Puro DEST (Addgene plasmid 17293), psPAX2 (Addgene plasmid 12260) and pMD2.G (Addgene plasmid 12259). The constructed vectors with their full sequence information are available in Professor Anurag Tandon’s lab of Department of Medicine, Tanz Centre for Research in Neurodegenerative Diseases, University of Toronto (Krembil Discovery Tower, 60 Leonard Avenue, 4th Floor-4KD481, Toronto, Ontario, Canada, M5T 2S8). These constructed vectors include pWPI/α-SynWt/Neo, pWPI/α-SynA30P/Neo, pWPI/α-SynA53T/Neo, pWPI/EGFP/Neo, pWPI/hPlk2Wt/Neo, pWPI/hPlk2K111M/Neo, pWPI/hPlk2T239D/Neo, pWPI/hPlk2T239V/Neo, pWPI/β5Wt/Neo, pWPI/β5T1A/Neo, pWPI/GDIWt/Neo, pWPI/GDIR218E/Neo, pWPI/GDIR240A/Neo, pWPI/Rab3AWt/Neo, pWPI/Rab3AT36N/Neo, pWPI/Rab3AQ81L/Neo, pLenti/TO/β5Wt/Puromycin (Puro), pLenti/TO/β5T1A/Puro.

## Author’s contributions

GZ and TW conceived the idea. GZ designed the experiments, performed the experiments and carried out data analysis. GZ and TW wrote the paper. All authors read and approved the final manuscript.

## Consent for publication

Not applicable

## Ethics approval and consent to participate

Not applicable

## References

1 Naldini L, Blomer U, Gallay P et al. In vivo gene delivery and stable tranpduction of nondividing cells by a lentiviral vector. Science 1996; 272: 263–67.

2 Cui Y, Golob J, Kelleher E et al. Targeting transgene expression to antigen-presenting cells derived from lentivirus-transduced engrafting human hematopoietic stem/progenitor cells. Blood 2002;99:399–408.

3 Lois C, Hong EJ, Pease S et al. Germline transmission and tissue-specific expression of transgenes delivered by lentiviral vectors. Science 2002;295:868–72.

4 Zufferey R, Dull T, Mandel RJ et al. Self-inactivating lentivirus vector for safe and efficient in vivo gene delivery. J Virol 1998;72:9873–80.

5 Hanna J, Markoulaki S, Schorderet P et al. Direct reprogramming of terminally differentiated mature B lymphocytes to pluripotency. Cell 2008;133:250–64.

6 Hotta A, Cheung AY, Farra N et al. Isolation of human iPS cells using EOS lentiviral vectors to select for pluripotency. Nat Methods 2009;6:370–76.

7 Sommer CA, Stadtfeld M, Murphy GJ et al. Induced pluripotent stem cell generation using a single lentiviral stem cell cassette. Stem Cells 2009;27:543–49.

8 Kabadi AM, Ousterout DG, Hilton IB et al. Multiplex CRISPR/Cas9-based genome engineering from a single lentiviral vector. Nucleic Acids Res 2014;42:e147.

9 Sanjana NE, Shalem O, Zhang F. 2014. Improved vectors and genome-wide libraries for CRISPR screening. Nat Methods 2014;11:783–84.

10 Shalem O, Sanjana NE, Hartenian E et al. Genome-scale CRISPR-Cas9 knockout screening in human cells. Science 2014;343:84–87.

11 Tiscornia G, Singer O, Verma IM. Production and purification of lentiviral vectors. Nat Protocols 2006;1:241–45.

12 Pfeifer A, Ikawa M, Dayn Y et al. Transgenesis by lentiviral vectors: lack of gene silencing in mammalian embryonic stem cells and preimplantation embryos. Proc Natl Acad Sci USA 2002;99:2140–45.

13 Geraerts M, Willems S, Baekelandt V et al. Comparison of lentiviral vector titration methods. BMC Biotech 2006;6:34.

14 Zhang G, Tandon A. Quantitative assessment on the cloning efficiencies of lentiviral transfer vectors with a unique clone site. Sci Rep 2012;2:415.

15 Zhang G, Tandon A. Combinatorial Strategy: A highly efficient method for cloning different vectors with various clone sites. Am J Biomed Res 2013;1:112–19.

16 Zhang G. A New Overview on the Old Topic: The Theoretical Analysis of “Combinatorial Strategy” for DNA Recombination. Am J Biomed Res 2013;1:108–11.

17 Zhang G, Zhang Y. On the “All or Half” Law of Recombinant DNA. Am J Biomed Res 2016;4:1–4.

18 Campeau E, Ruhl VE, Rodier F et al. A versatile viral system for expression and depletion of proteins in mammalian cells. PLoS One 2009;4: e6529.

19 Visanji NP, Wislet-Gendebien S, Oschipok LW et al. The effect of S129 phosphorylation on the interaction of α‐synuclein with synaptic and cellular membranes. J Biol Chem 2011;286:35863–73.

20 Kanemoto S, Griffin J, Markham-Coultes K et al. Proliferation, differentiation and amyloid-β production in neural progenitor cells isolated from TgCRND8 mice. Neuroscience 2014;261:52–59.

21 Qiu WQ, Walsh DM, Ye Z et al. Insulin-degrading enzyme regulates extracellular levels of amyloid – protein by degradation. J Biol Chem 1998;273:32730–38.

22 Chapman JM, Knoepp SM, Byeon MK et al. The colon anion transporter, down-regulated in adenoma, induces growth suppression that is abrogated by E1A. Cancer Res 2002;62:5083–88.

23 Kobayashi T, Wang T, Maezawa M et al. Overexpression of the oncoprotein prothymosin-α triggers a p53 response that involves p53 acetylation. Cancer Res 2006;66:3137–44.

24 Yang L, Guell M, Byrne S et al. Optimization of scarless human stem cell genome editing. Nucleic Acids Res 2013;41:9049–61.

25 Dekker LV, Daniels Z, Hick C et al. Analysis of human Na_v_1.8 expressed in SH-SY5Y neuroblastoma cells. Eur J Pharmacol 2005;528:52–58.

26 Miyagi S, Nishimoto M, Saito T et al. The *Sox2* regulatory region 2 functions as a neural stem cell-specific enhancer in the telencephalon. J Biol Chem 2006;281:13374–81.

27 Corti S, Nizzardo M, Nardini M et al. Embryonic stem cell-derived neural stem cells improve spinal muscular atrophy phenotype in mice. Brain 2010;133:465–81.

28 Lee H, Cha S, Lee MS et al. Role of antiproliferative B cell translocation gene-1 as an apoptotic sensitizer in activation-induced cell death of brain microglia. J Immunol 2003;171:5802–11.

29 Schmidtke G, Kraft R, Kostka S et al. Analysis of mammalian 20S proteasome biogenesis: the maturation of betasubunits is an ordered two-step mechanism involving autocatalysis. EMBO J 1996;15:6887–98.

30 Li Z, Arnaud L, Rockwell P et al. A single amino acid substitution in a proteasome subunit triggers aggregation of ubiquitinated proteins in stressed neuronal cells. J Neurochem 2004;90:19–28.

31 Chen RHC, Wislet-Gendebien S, Samuel F et al. α-Synuclein membrane association is regulated by the Rab3a recycling machinery and presynaptic activity. J Biol Chem 2013;288:7438–49.

32 Sastry L, Johnson T, Hobson MJ et al. Titering lentiviral vectors: comparison of DNA, RNA and marker expression methods. Gene Ther 2002;9:1155–62.

33 Miyoshi H, Blomer U, Takahashi M et al. Development of a self-inactivating lentivirus vector. J Virol 1998;72:8150–57.

34 Chalfie M, Tu Y, Euskirchen G et al. Green fluorescent protein as a marker for gene expression. Science 1994;263:802–05.

35 Sambrook J, Russell DW. Molecular Cloning: A Laboratory Manual. 2001; Cold Spring Harbor Laboratory Press: Plainview, NY.

36 Yamada K, Olsen JC, Patel M et al. Functional correction of fanconi anemia group C hematopoietic cells by the use of a novel lentiviral vector. Mol Ther 2001;3:485–90.

37 Davis JL, Witt RM, Gross PR et al. Retroviral particles produced from a stable human-derived packaging cell line transduce target cells with very high efficiencies. Hum Gene Ther 1997;8:1459–67.

38 Kirkwood TBL, Bangham CRM. Cycles, chaos, and evolution in virus cultures: a model of defective interfering particles. Proc Natl Acad Sci USA 1994;91:8685–89.

39 Doux JML, Morgan JR, Snow RG et al. Proteoglycans secreted by packaging cell lines inhibit retrovirus infection. J Virol 1996;70:6468–73.

40 Naldini L, Blomer U, Gage FH et al. Efficient transfer, integration, and sustained long-term expression of the transgene in adult rat brains injected with a lentiviral vector. Proc Natl Acad Sci USA 1996;93:11382–88.

